# Segmentation of the Zebrafish Brain Vasculature from Light Sheet Fluorescence Microscopy Datasets

**DOI:** 10.1101/2020.07.21.213843

**Authors:** Elisabeth C. Kugler, Andrik Rampun, Timothy J.A. Chico, Paul A. Armitage

## Abstract

Light sheet fluorescent microscopy allows imaging of zebrafish vascular development in great detail. However, interpretation of data often relies on visual assessment and approaches to validate image analysis steps are broadly lacking. Here, we compare different enhancement and segmentation approaches to extract the zebrafish cerebral vasculature, provide comprehensive validation, study segmentation robustness, examine sensitivity, apply the validated method to quantify embryonic cerebrovascular volume, and examine applicability to different transgenic reporter lines. The best performing segmentation method was used to train different deep learning networks for segmentation. We found that U-Net based architectures outperform SegNet. While there was a slight overestimation of vascular volume using the U-Net methodologies, variances were low, suggesting that sensitivity to biological changes would still be obtained.

**Highlights:**

- General filtering is less applicable than Sato enhancement to enhance zebrafish cerebral vessels.
- Biological data sets help to overcome the lack of segmentation gold-standards and phantom models.
- Sato enhancement followed by Otsu thresholding is highly accurate, robust, and sensitive.
- Direct generalization of the segmentation approach to transgenics, other than the one optimized for, should be treated with caution.
- Deep learning based segmentation is applicable to the zebrafish cerebral vasculature, with U-Net based architectures outperforming SegNet architectures.

**Graphical Abstract:** 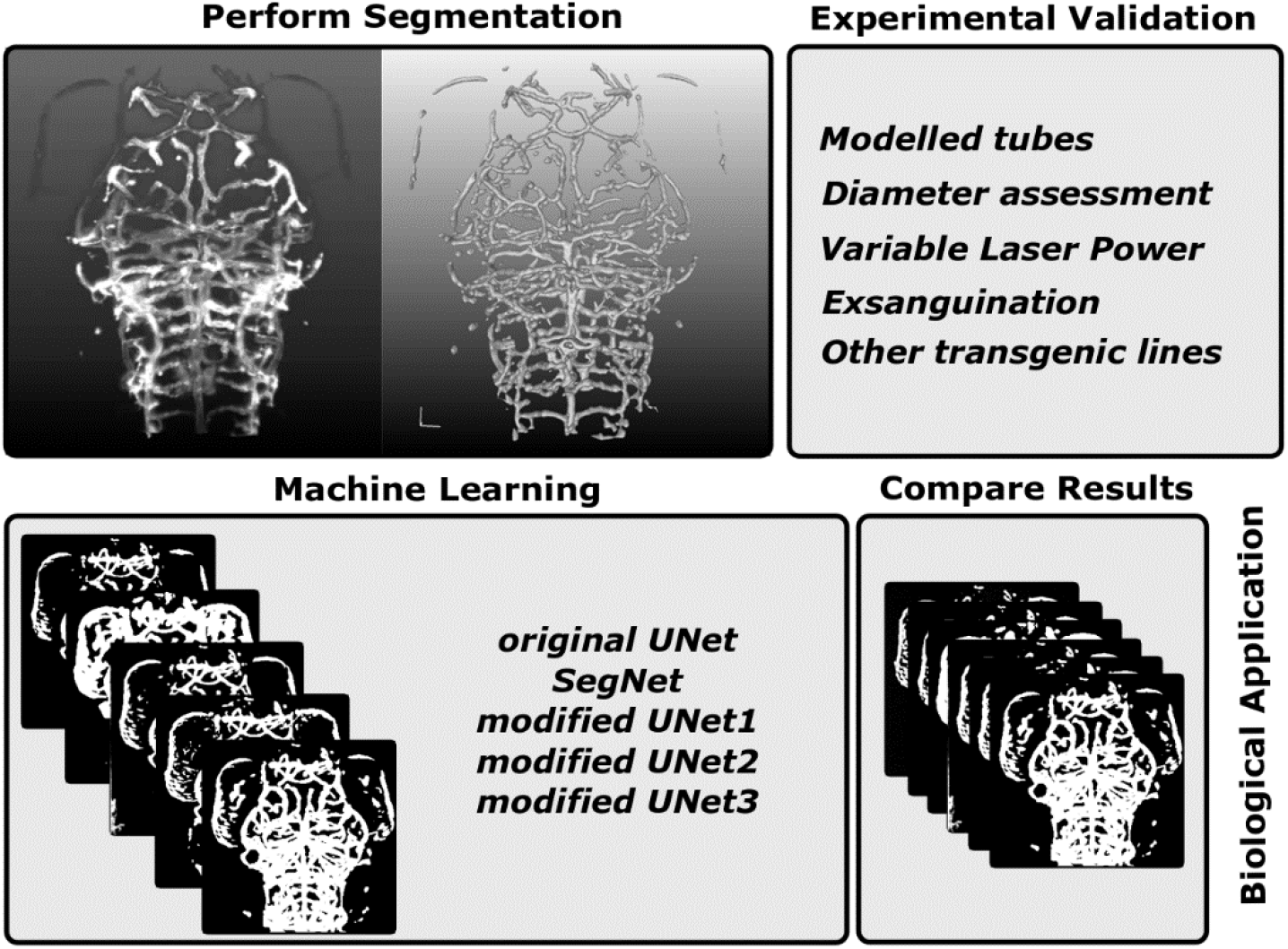

## 1. Introduction

### 1.1. Zebrafish in Cardiovascular Research

Vascular diseases are the leading cause of death worldwide (Feigin Valery L. et al., 2017; Lackland and Weber, 2015), with diseases of the central nervous system associated with neurodegeneration, arteriovenous malformations, aneurysms and stroke.

Zebrafish are used to study vascular development and disease due to characteristics including high genomic similarity to human, fecundity, rapid *ex utero* development, and larval transparency (Bakkers, 2011; Chico et al., 2008; Gut et al., 2017). The availability of fluorescent transgenic reporter lines allows visualization of sub-cellular structures of interest with high specificity. For example, endothelial cells which line the vascular lumen can be visualized non-invasively and *in vivo*, replacing laborious microangiography (Lawson and Weinstein, 2002).

With the emergence of sophisticated microscopy techniques, such as light sheet fluorescence microscopy (LSFM), it is possible to acquire vascular information in greater anatomical detail and over prolonged periods of time (hours to days) (Huisken et al., 2004).

Together, transgenic lines and LSFM allow data acquisition that is rich in anatomical depth, spatiotemporal resolution, and detail, meaning that the limitation of experimental throughput and assessment has now become data handling and analysis, rather than data acquisition.

### 1.2. Challenges in quantifying the zebrafish cranial vasculature

While some aspects of the zebrafish vascular architecture are visually apparent without quantification (such as increasing network complexity during development), others may be too subtle for human perception (such as diameter changes). Computational quantification of the vascular architecture in 3D is not just more comprehensive (e.g. providing measurements of vessel diameter, length, branching, etc.), but also reproducible (e.g. overcoming subjective bias or inter-observer variability) than human assessment. However, while quantification of vascular geometry is an active research field in the medical domain, it has received less attention in pre-clinical models such as zebrafish.

The main reasons for the lack of a robust segmentation and quantification approach for the zebrafish cerebral vasculature are: **(i)** The majority of zebrafish vascular research has focused on vascular development in the trunk, as trunk vessel formation shows a highly stereotypic growth pattern that is well characterised. **(ii)** The zebrafish cerebral vasculature is increasingly studied, but is topologically more complex than the trunk vasculature, presenting significant challenges to overcome. **(iii)** As endothelial cells are visualized in transgenic lines, lumenized vessels display a cross-sectional double-peak intensity distribution, while small / unlumenized vessels have a single-peak distribution, requiring image analysis approaches that detect and discriminate between these cases. **(iv)** LSFM is a relatively new technique with commercial microscopes only being available in recent years. **(v)** Zebrafish are still emerging as a model and have not received the same level of attention as other models, such as rodent. **(vi)** Data acquisition with light sheet fluorescence microscopy produces large datasets, requiring more computational resources for storage and processing.

Additionally, due to the lack of gold-standards or phantoms for image processing and analysis, developed analysis approaches are rarely examined in-depth and it is unclear how accurate or sensitive the suggested methods are.

### 1.3. Previous work aiming to quantify the zebrafish cerebral vasculature

Quantification of left hind-brain vessels was previously performed by Tam et al. (2012), while Chen et al. (2012) presented quantification of the mid-brain vasculature. Both methods used confocal microscopy and focused only on a sub-region of the cerebral vasculature rather than the whole cerebral vasculature.

In the study of Tam et al. (2012), measurement of vascular density and diameter were performed after deconvolution using Imaris. Chen et al. (2012) quantified vessel length, branching hierarchy (Strahler, 1952), existence of loops, and vascular pruning events using the commercial software Neurolucida. However, neither approach provided sufficient methodological detail to facilitate replication and perform an in-depth assessment of their performance.

Recently, a segmentation approach using machine learning was suggested for the whole zebrafish embryonic vasculature in LSFM data (Daetwyler et al., 2019). The method was trained on data from double-transgenic zebrafish providing endothelial as well as luminal signal. As additional luminal signal is required to extract vascular information, data-load is doubled compared to a single transgenic and only visual assessment of segmentation outcomes was performed, with no further validation. Previously, Kugler *et al.* presented methods to enhance the cerebral vasculature in LSFM data using general filtering (GF; Median Filter and Rolling ball) (Kugler et al., 2018) and enhancement utilizing the Hessian matrix with the assumption of local vessel tubularity, based on the filter proposed by Sato et al. (Kugler et al., 2019; Sato et al., 1997). This was further complemented by investigation of different segmentation approaches, which were readily implemented in the Fiji image analysis framework (Schindelin et al., 2012), but no validation of the suggested approach was performed. For zebrafish cerebral vascular quantification, no ground truth exists as there has been no previous attempt to assess segmentation quality.

We here aim to overcome the lack of knowledge about segmentation quality by performing detailed validation of our segmentation methodology using simulated data, manual measurements, and various *in vivo* biological datasets that challenge segmentation performance in different ways. Thus, address the lack of a gold-standard experimentally and analytically.

### 1.4. Contributions of this work

Our study makes the following specific contributions (**Fig. 1**):

**Figure 1.**
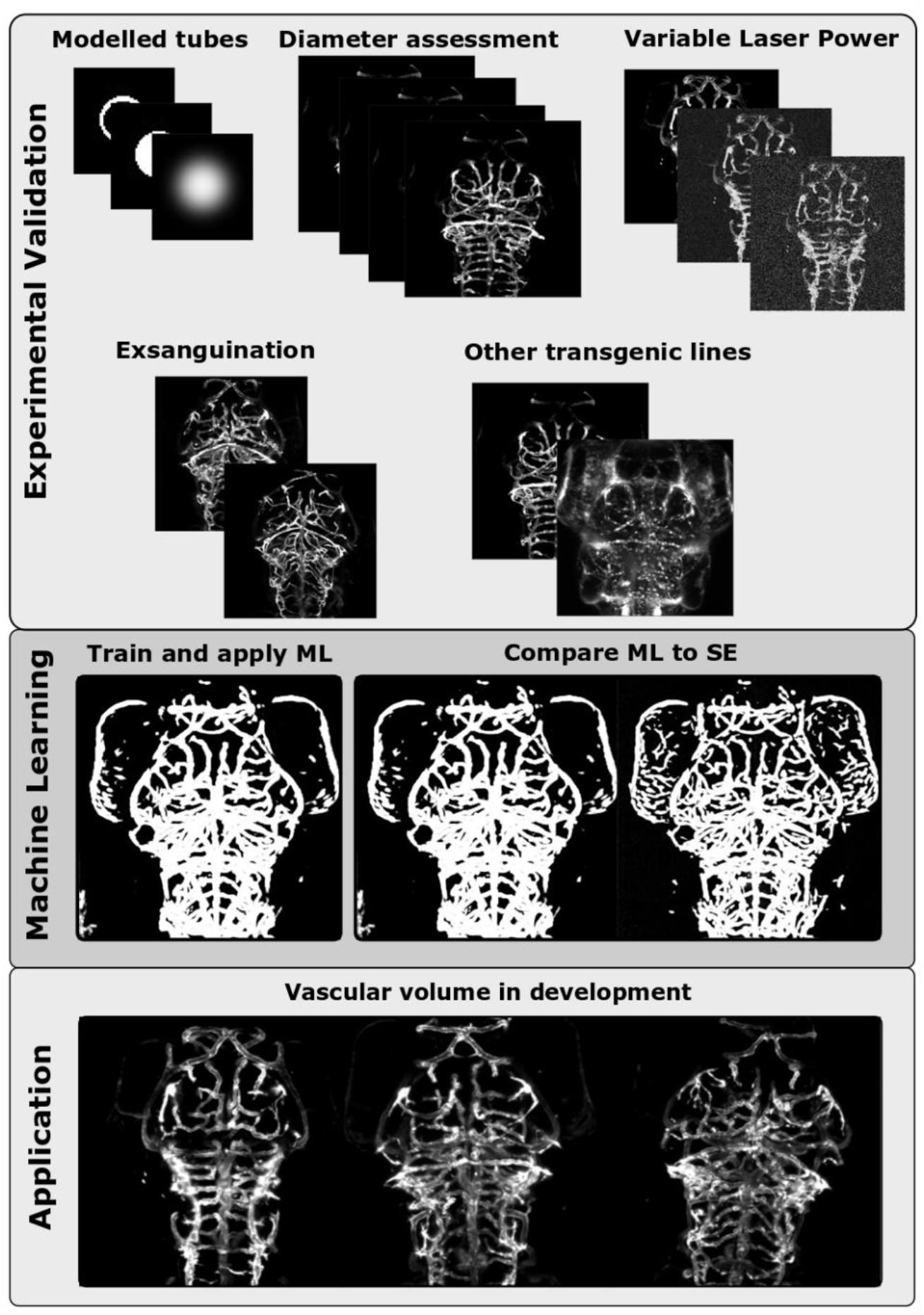
Workflow of data analysis presented in this paper.

We analyse the impact of different modelled tubes, with simulated parameters chosen to be realistic for the zebrafish vasculature, on segmentation outcomes following Sato enhancement (SE) applied at different scales (Sato et al., 1997).

Given that there is no gold-standard for zebrafish vascular segmentation, we present several methodological approaches to allow for the assessment of segmentation robustness, sensitivity, and accuracy. We addressed this by establishing datasets that challenge the segmentation performance to understand whether successful segmentation would be achieved and thus true biological vascular volumes could be extracted. These examined datasets were composed as follows:

i. To assess segmentation accuracy, detailed assessment of vessel diameters obtained from manual measurements are compared to those obtained after enhancement and segmentation.
ii. To assess noise sensitivity, we examine a dataset with a controlled decrease of image quality, as quantified by contrast-to-noise ratio (CNR), via data augmentation and reduction of laser power during repeated image acquisition.
iii. To assess segmentation sensitivity to true biological changes, *i.e.* blood loss resulting in vascular volume decrease, segmentation is performed on data acquired prior to and after exsanguination.
iv. Following segmentation validation we present the application of our suggested enhancement and segmentation approach to quantify the cerebral vascular volume from 3-to-5 days post fertilization (dpf).
v. To study applicability to other transgenic lines, vascular segmentation is performed in the double-transgenic lines *Tg(fli1a:eGFP)*^*y1*^, *Tg(kdrl:HRAS-mCherry)*^*s916*^ (Chi et al., 2008; Lawson and Weinstein, 2002), *Tg(fli1a:CAAX-eGFP), Tg(kdrl:HRAS-mCherry)*^*s916*^ (Gebala et al., 2016), as well as *Tg(fli1a:LifeAct-mClover)*^*sh467*^, *Tg(kdrl:HRAS-mCherry)*^*s916*^ (Savage et al., 2019). *Tg(kdrl:HRAS-mCherry)*^*s916*^ has a high CNR (Kugler et al., 2019) and a vascular specific expression pattern than the other examined transgenics, which are also commonly used in zebrafish cardiovascular research labs worldwide. Lastly, we use our validated segmentation approach based on SE to generate data used to train the original U-Net (Ronneberger et al., 2015), SegNet (Badrinarayanan et al., 2017), and three modified versions of the original U-Net architecture (dU-Net) and analyse their performance.

## 2. Material and Methods

### 2.1. Zebrafish Husbandry

Experiments were performed according to the rules and guidelines of institutional and UK Home Office regulations under the Home Office Project Licence 70/8588 held by TC. Maintenance of adult zebrafish *Tg(kdrl:HRAS-mCherry)*^*s916*^ (Chi et al., 2008) and *Tg(fli1a:eGFP)*^*y1*^ (Lawson and Weinstein, 2002) was performed as described in standard husbandry protocols (Westerfield, 1993). Embryos, obtained from controlled mating, were kept in E3 medium buffer with methylene blue and staged as previously described Kimmel et al. (Kimmel et al., 1995).

### 2.1. Image Acquisition Settings and Properties

Anaesthetized embryos were embedded in 2% LMP-agarose with 0.01% Tricaine in E3 (MS-222, Sigma). Data acquisition of the cerebral vasculature was performed using a Zeiss Z.1 light sheet microscope, Plan-Apochromat 20x/1.0 Corr nd=1.38 objective, dual-side illumination with online fusion, activated Pivot Scan, image acquisition chamber incubation at 28°C, with a scientific complementary metal-oxide semiconductor (sCMOS) detection unit. The properties of acquired data were as follows: 16bit image depth, 1920 × 1920 × 400-600 voxel (x,y,z; approximate voxel size of 0.33 × 0.33 × 0.5 μm, respectively). Multicolour images in double-transgenic embryos were acquired in sequential mode.

### 2.2. Datasets

#### 2.2.1. Modelled Tubes

Modelled straight tubes (hollow, filled and Gaussian blurred) were simulated with a uniform signal intensity of 255 against zero background intensity using Fiji (Schindelin et al., 2012). Total image size was 268 × 268 × 250 voxels with voxel size 0.2 × 0.2 × 0.33 μm. Tubes were produced to resemble the following biological settings: (i) hollow tubes – 20 μm outer diameter (1.13 μm wall thickness) and 8.3 μm outer diameter (0.8 μm wall thickness); resembling lumenized vessels; (ii) filled tubes - resembling unlumenized vessels (with the same outer diameter as above); (iii) Gaussian blurred tubes were created by taking the simulated datasets in (i) and (ii) and applying a Gaussian filter with a sigma of 5 voxels to resemble a more realistic intensity distribution of fluorescence; (iv) increasing Gaussian white noise with standard deviation 25, 50 or 100 (zero background intensity), resembling background noise was added to the simulated datasets produced in (i), (ii) and (iii). The modelled tubes were used to establish the Sato filter response for different types of tubes when the filter is applied with varying scale parameters. We were particularly interested in establishing how the filter responds to a double-peak distribution from a hollow tube, as typically found in lumenized vessels.

#### 2.2.2. Transgenic Zebrafish

Data to test vascular enhancement approaches were acquired at 4 days post fertilization (dpf). Data analysed for assessment of segmentation robustness included the following:

i. Dataset with controlled decrease of vascular contrast-to-noise ratio (CNR) by decrease of laser power during repeated acquisition (Kugler et al., 2018) (laser power (LP) 1.2%, 0.8% and 0.4%; exposure 30ms for all; 4dpf; n=10 embryos from 2 experimental repeats). Augmented data were produced from LP 1.2% by addition of Gaussian noise with mean of zero and standard deviation of 50 using Fiji (Schindelin et al., 2012).
ii. Data acquisition before and after exsanguination by mechanical opening of the heart cavity with forceps in *Tg(kdrl:HRAS-mCherry)*^*s916*^ (4dpf; n=16 embryos from 2 experimental repeats).
iii. Double-transgenics: *Tg(fli1a:eGFP)*^*y1*^, *Tg(kdrl:HRAS-mCherry)*^*s916*^ (n=21 embryos from 2 experimental repeats), *Tg(fli1a:CAAX-eGFP), Tg(kdrl:HRAS-mCherry)*^*s916*^ (n=17 embryos from 2 experimental repeats), as well as *Tg(fli1a:LifeAct-mClover)*^*sh467*^, *Tg(kdrl:HRAS-mCherry)*^*s916*^ (n=23 embryos from 3 experimental repeats).
iv. Data to quantify vascular volume during early development were acquired in *Tg(kdrl:HRAS-mCherry)*^*s916*^ (3dpf n=12, 4dpf n=13, 5dpf n=15; all data from 2 experimental repeats).
v. In order to train the deep learning networks, a *Tg(kdrl:HRAS-mCherry)*^*s916*^ training dataset was acquired at 4dpf n=9 and a separate evaluation dataset was acquired to evaluate the performance of the network using the same age and transgenic line with n=10.

### 2.3. Image Analysis

#### 2.3.1. Basic Analysis

All image analysis, pre-processing and segmentation were performed using the open-source software Fiji (Schindelin et al., 2012). Contrast-to-noise ratio (CNR) was quantified and motion correction performed as described in (Kugler et al., 2019). Full-Width-Half-Maximum (FWHM; *Eq. 1*) was calculated from extracted cross-sectional line region of interest (ROI) intensity profiles, *f*(*x*), using Matlab.

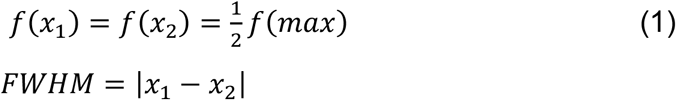

#### 2.3.2. Image Enhancement

The following vascular enhancement methods were studied:

i. General filtering (GF): 2D median filter with a radius of 6 voxels (13-by-13 neighbourhood) (Lim, 1990) and rolling ball algorithm of size 200 (Sternberg, 1983), as presented in (Kugler et al., 2019).
ii. Sato Enhancement (SE): Enhancement based on the line enhancement filter proposed by Sato et al. (Sato et al., 1997) with the assumption of local vessel tubularity, as implemented into the Fiji Tubeness Plugin by Mark Longair, Stephan Preibisch and Johannes Schindelin (Schindelin et al., 2012), was applied as described in (Kugler et al., 2019).

#### 2.3.3. Segmentation and Total Vascular Volume Measurement

Segmentation of enhanced images was performed using global Otsu thresholding (Otsu, 1979) to distinguish vascular from non-vascular information. Following segmentation, the total dorsal cerebral vascular volume was quantified in a cerebral ROI defined as described in (Kugler et al., 2018).

#### 2.3.4. Deep Learning Architectures

We chose two popular deep learning network architectures, namely the original U-Net (Ronneberger et al., 2015) and SegNet (Badrinarayanan et al., 2017), and also modified the original U-Net architecture to suit our segmentation problem (called dU-Net). This was achieved by making the original U-Net architecture deeper by adding more convolutional layers, employing batch normalization, and applying dropout procedures to avoid overfitting (**Fig. 2**).

**Figure 2.**
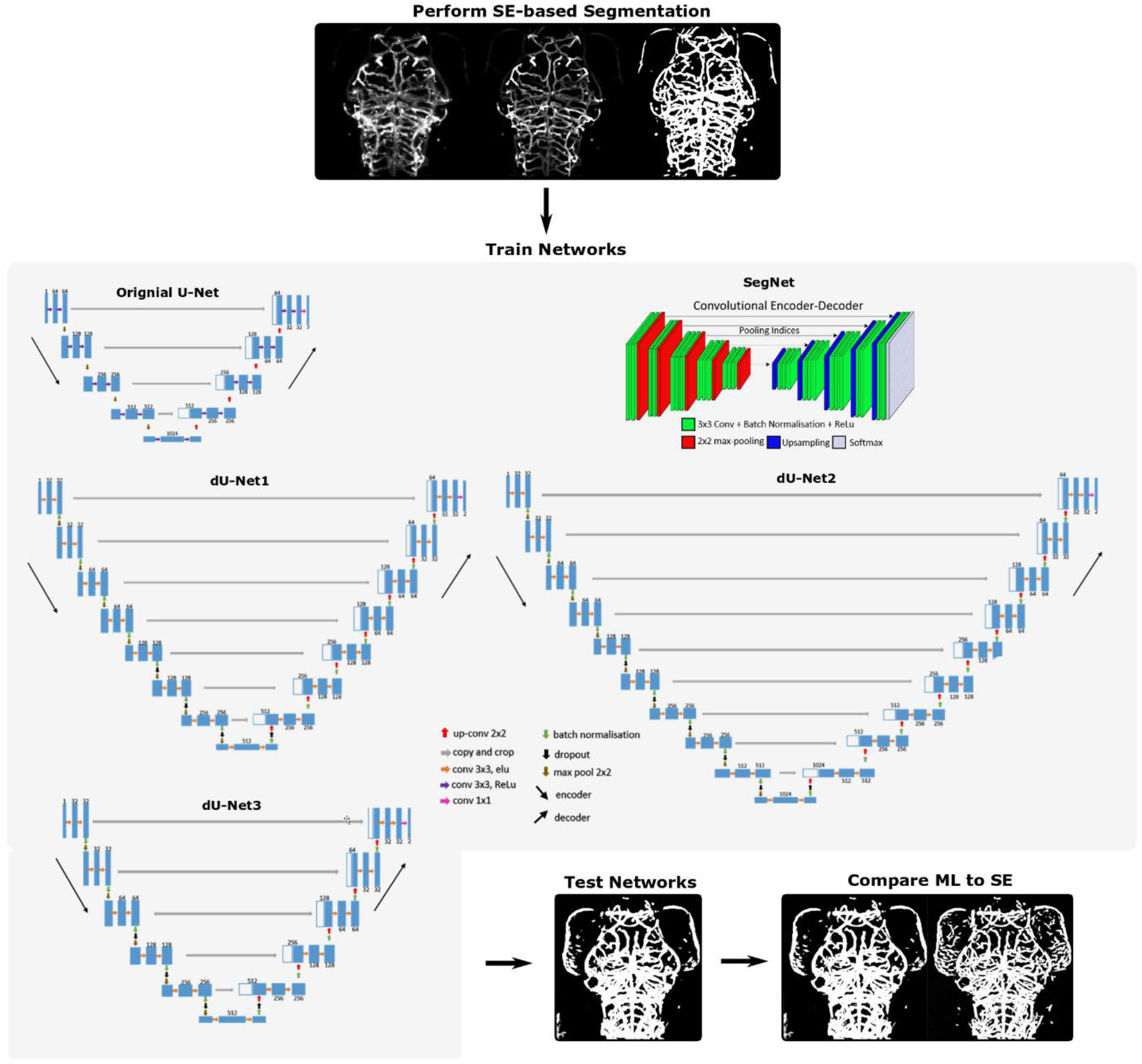
Machine learning workflow and network.

The U-Net architecture was developed specifically for segmenting biomedical images, whereas the SegNet architecture was developed for generic image segmentation. Both networks have shown promising results in many applications of semantic segmentation. However, employing the network architecture directly is often insufficient to produce good segmentation results. Therefore, we modified the original U-Net architecture to suit our segmentation problem.

The original U-net contains four levels of 3×3 convolutional layers in both encoder and decoder parts with up to 1024 feature channels used. Our first modification (dU-Net1), contains seven levels of 3×3 convolutional layers with a maximum of 512 feature channels used. We further modified the network by adding more convolutional layers, hence making it deeper, with nine levels of 3×3 convolutional layers with up to 1024 feature channels used (dU-Net2). Finally, the dU-Net3 has five levels of 3×3 convolutional layers with up to 512 feature channels. Note that the 3×3 convolutional operation with Rectified Linear Unit (ReLu) activation function is used in the original U-Net, whereas, in our modified versions we employ the Exponential Linear Unit (ELU) function instead. The ReLu activation function will output the summed weighted input directly if it is positive, otherwise, it will output zero. On the other hand, the ELU activation function uses a parameterized exponential function to transition from the positive to small negative values which are closer to zero. Mean activations that are closer to zero enable faster learning as they bring the gradient closer to the natural gradient (Clevert et al., 2016). In our previous work, we found that the ELU activation function provides better overall segmentation results compared to the ReLU (Rampun et al., 2020). Furthermore, a batch normalization function is added after each level convolutional operation in the dU-Net and as the number of feature channels increase, we employ the dropout function to avoid overfitting. **Table 1** summarizes the network architecture details. The dU-Net architectures were motivated by the following:

**Table 1.** Network Architecture Summary. Comparison between the original SegNet, U-Net and its modifications employed in this study.

|  | U-Net | SegNet | dU-Net1 | dU-Net2 | dU-net3 |
| --- | --- | --- | --- | --- | --- |
| **Depth (level)** | 4 | 5 | 7 | 9 | 5 |
| **Activation function** | ReLu | ReLu | ELU | ELU | ELU |
| **Kernel size** | 3x3 | 3x3 | 3x3 | 3x3 | 3x3 |
| **Batch Normalization** | None | Yes | Yes | Yes | Yes |
| **Maximum number of feature channels** | 1024 | 512 | 512 | 1024 | 512 |
| **Dropout** | None | None | 0.4 | 0.6 | 0.3 |
| **Loss function** | Pixel-wise Softmax with cross entropy | Binary cross entropy | Binary cross entropy+ Dice coefficient | Binary cross entropy+ Dice coefficient | Binary cross entropy+ Dice coefficient |

- **Data sparsity**. Only about 15% of pixels belong to the zebrafish cerebral vasculature (foreground), with 1-5% foreground pixels in bottom and top slices. Therefore, a balanced loss function is essential to handle the class imbalance problem (e.g. the imbalance in the number of pixels between foreground and background). Moreover, deeper network architectures can capture diverse coarse and finer features, allowing the extraction of more discriminant features which may not be available in a shallow network (e.g. the original U-Net).
- **Background fluorescence**. Non-specific signal (e.g. background and sample auto-fluorescence) in our data can be very noisy and appear similar to the cerebral vasculature. This can easily deteriorate the final segmentation results.

#### 2.3.5. Deep Learning Training, Validation, and Testing

All images were automatically down-sampled to 256×256 to decrease the memory and time required for training. The following data augmentations were applied: **(i)** random rotation range of up to 180°, **(ii)** zooming in and out with a range of 0:1 to 2:0, and **(iii)** horizontal and vertical flips.

To assess the performance, the networks were trained on a dataset of *Tg(kdrl:HRAS-mCherry)*^*s916*^ and applied to another (4dpf n=10).

Using the training dataset, we trained our network with the Root Mean Square Propagation (RMSprop) (Hinton et al.) implementation in Keras with a Tensorflow back-end. The initial learning rate (lr) and gradient moving average decay factor were 0.0003 and 0.8, respectively. To maximize the network speed and to have a better estimation of the gradient of the full dataset (hence faster convergence), we favoured a maximum batch size (bs = 32).

The number of iterations used per epoch (E) was based on the number of samples divided by batch size. We monitored the Dice and Jaccard coefficients, set E = 1000 and applied the ‘*EarlyStopping*’ mechanism on the validation set to automatically stop training when the loss function value did not change after 50 epochs. To minimise covariance shifts, for each convolutional block we applied the batch normalisation procedure, which further increased the learning process. The loss function was computed as a combination of binary cross-entropy (our study can be seen as a binary classification: vascular versus non-vascular pixels) and Dice Coefficient, which is described as:

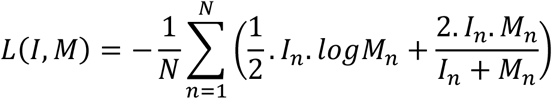

where *I*_*n*_ and *M*_*n*_ are the 2D training images and their corresponding 2D ground truth binary image, respectively.

For weight initialization, we followed the weight initialisation procedure implemented in the original paper (Ronneberger et al., 2015) using the Gaussian distribution with a standard deviation of 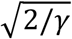, where *γ* is the number of incoming nodes of one neuron.

#### 2.3.6. Statistics and Data Representation

Gaussian distribution conformation was evaluated using the D’Agostino-Pearson omnibus test [21]. Statistical analysis was performed using One-way ANOVA or paired Students t-test in GraphPad Prism Version 7 (GraphPad Software, La Jolla California USA). Statistical significance was represented as: p<0.05 *, p<0.01 **, p<0.001 ***, p<0.0001 ****. Graphs show mean values ± standard deviation unless otherwise indicated. Image representation and visualization was done with Inkscape Version 0.48 (https://www.inkscape.org). Bland-Altman measurement was used to assess segmentation methods: plotting the ratio (SE volume / DL volume) versus average of both volume measurements. Images were visualized as maximum intensity projections (MIPs) using false-colour representation and intensity inversion where appropriate.

## 3. Results and Discussion

### 3.1. Vessel enhancement based on vascular tubularity and the impact of input shape

To test vessel enhancement using a shape-based filter with the assumption of local vascular tubularity, the Sato enhancement (SE) filter (Sato et al., 1997) already implemented into Fiji was evaluated for its applicability to images of the zebrafish vasculature acquired with LSFM. Filters were applied at a scale size similar to the average vessel size of cerebral vessels under the assumption that this would return an optimum response (scale 10μm).

Comparing three vessels in original *in vivo* data (**Fig. 3A**) before and after SE showed satisfactory vessel enhancement for all vessels examined (**Fig. 3A’**). We further examined whether similar results would be obtained with inverted intensity (dark vessels on bright background; **Fig. 3B**), as SE is theoretically unable to perform in this state. Indeed, SE was unable to enhance vessels in this dark-on-bright state (**Fig. 3B’**). Overall, SE delivered meaningful results provided that data is presented as bright-on-dark.

**Figure 3.**
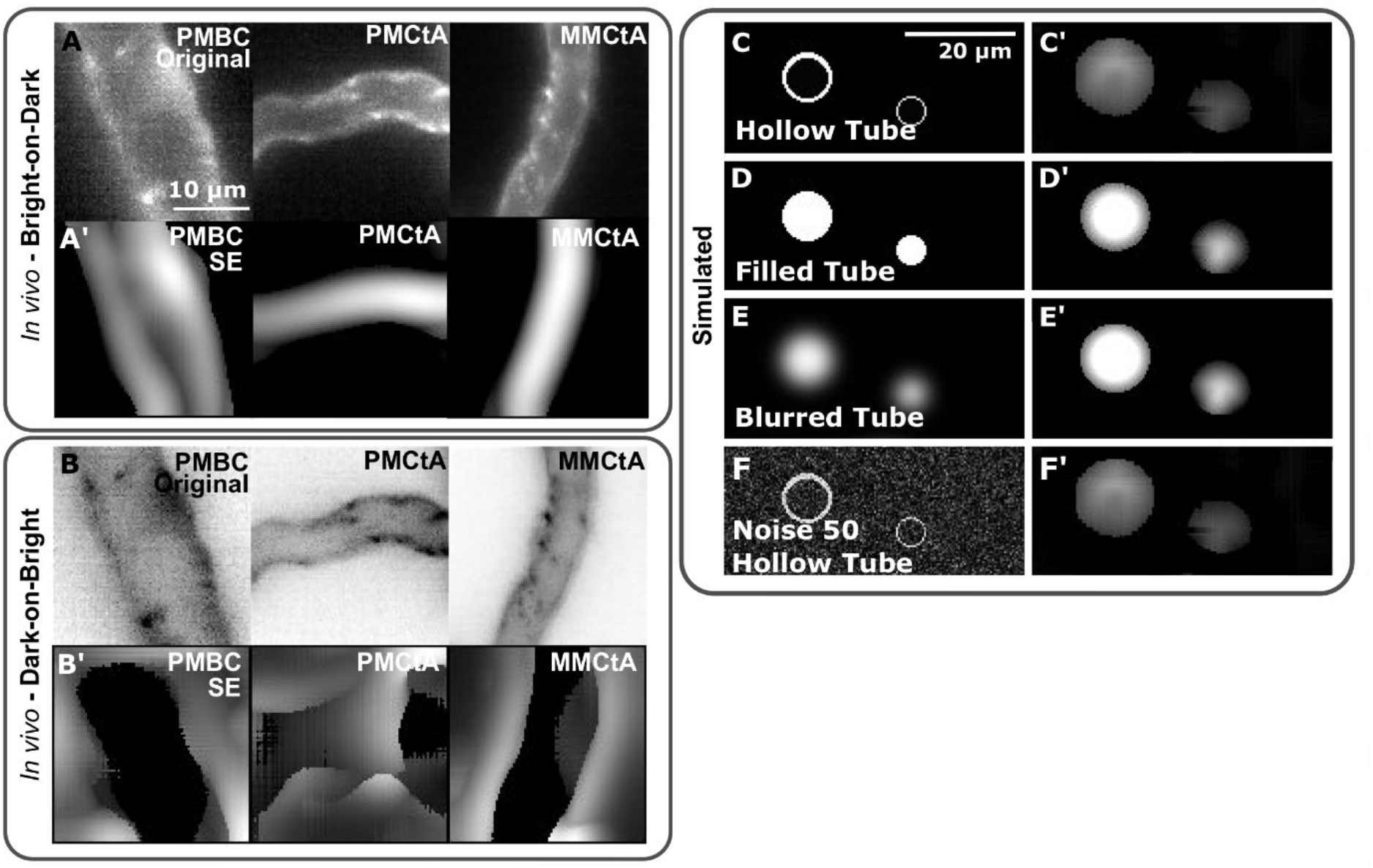
Vessel enhancement based on vascular tubularity and the impact of input shape. **(A)** Original data with bright vessels on dark background. **(A’)** Data processed with Sato enhancement showed successful enhancement. **(B)** Inverted data with dark vessels on bright background. **(B’)** SE was not able to perform after greyscale inversion. **(C,C’)** Applying SE to hollow tubes results in double-to-single peak conversion. **(D,D’)** Enhancement of filled tubes with SE results in successful enhancement. **(E,E’)** SE enhancement results for Gaussian blurred tubes were similar to unblurred tubes. **(F,F’)** Addition of artificial Gaussian noise at level 50 did not significantly alter SE enhancement results.

We further wanted to assess whether input shape would impact the filter output after SE and whether lumenized (hollow tubes / double-peak intensity) and unlumenized / unperfused (filled tubes / single intensity peak) vessels would need to be considered separately. This was of interest as SE was originally developed for magnetic resonance imaging (MRI) data which display a single-peak intensity distribution that can be approximated by a Gaussian distribution.

Applying SE to the simulated hollow tubes converted these to filled tubes (double-to-single peak conversion) when enhancement scale was approximately at the size of tubes (10μm; **Fig. 3C**). Enhancement of filled and filled Gaussian blurred tubes was similar (**Fig. 3D,E**). Addition of artificial Gaussian noise did not significantly alter enhancement (**Fig. 3F**).

These data showed that SE was able to convert double-to-single peak intensity distributions if applied at the scale of tubes, suggesting that lumenized and unlumenized vessels would both be enhanced similarly. Importantly, blurred tubes and data with additional noise were also enhanced successfully, suggesting that SE should be applicable to typical zebrafish data acquired with LSFM. However, there was a tendency for the simulated tube width to appear broader after enhancement, so any subsequent segmentation would need to be tuned to recover the correct vessel width.

### 3.2. Segmentation accuracy

To evaluate segmentation *accuracy*, vessel diameters obtained after applying the proposed enhancement and segmentation methodologies were compared to those obtained by “gold standard” manual measurement. This allows us to identify whether our segmentation approach will deliver true biological results across the expected range of vessel sizes, or if any systematic errors arise. Manual measurements of four vessels spanning a range of diameters were obtained; central artery (CtA) with an average diameter of 8.154±1.27μm, middle mesencephalic central artery (MMCtA) with a diameter of 9.78±2.09μm, primordial midbrain channel (PMBC) with a diameter of 11.14±1.68μm, basilar artery (BA) with a diameter of 22.28±3.89 μm (n=12 3dpf embryos).

As it is time consuming and prone to observer bias to perform manual vessel diameter measurement, we investigated whether full-width-half-maximum (FWHM) of the vessel cross-sectional intensity profile would provide a good estimate of vessel diameter. When comparing FWHM with the manual diameter measurements, no statistically significant differences were found (**Fig. 4A**; all vessels p>0.9999) and measurement errors were small (CtA 1.44μm, MMCtA 0.91μm, PMBC 1.35μm, BA 3.83μm). Assessment of Pearson Correlation and Bland-Altman analyses showed no systematic error, suggesting that the observed differences were unbiased. However, three outliers were observed in the FWHM measurements where poor agreement with the manual measurements was observed (**Fig. 4A**; unfilled arrowhead). These were found to be caused by strong asymmetric cross-sectional double-peak intensity distributions (**Fig. 4B**). We anticipate that this could be overcome by averaging cross-sectional intensity profiles from several different orientations to produce an average diameter for the vessel of interest. Thus, we suggest that the FWHM can be used to measure vascular diameter in zebrafish reporter lines with the *caveat* that outliers need to be considered.

**Figure 4.**
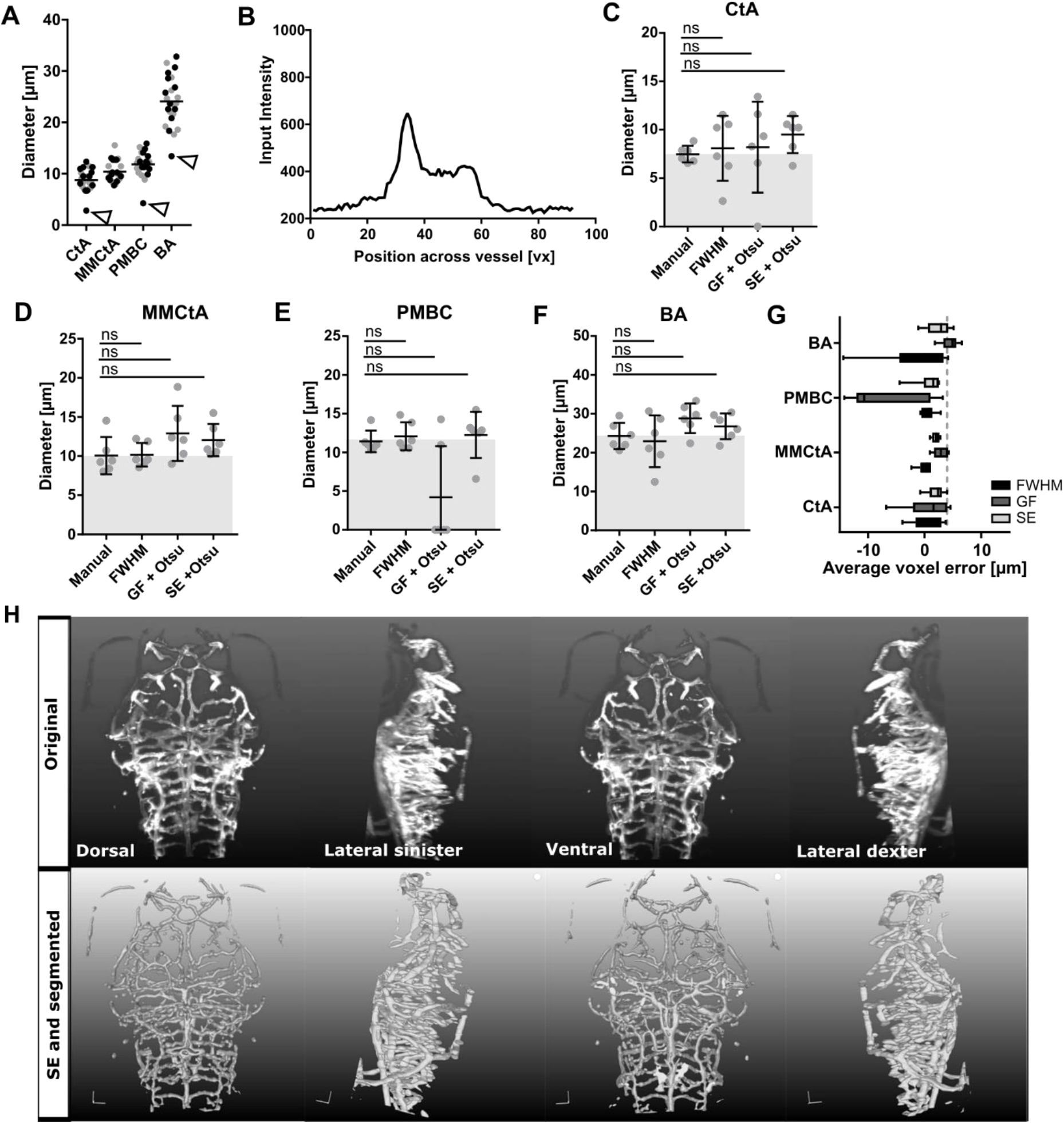
Validation of segmentation accuracy. **(A)** Comparing manual measurements (grey dots) to automated FWHM (black dots) showed a good agreement. White arrowheads indicate outliers which were caused by skewed cross-sectional intensity distributions (**B**). Comparison of manual measurements to FWHM, after GF with thresholding and SE with thresholding in the CtA **(C)**, MMCtA **(D)**, PMBC **(E)**, and BA **(F**; C-F Kruskal-Wallis test). **(G)** Average voxel error is independent of vessel diameter (averaged n=6 3dpf embryos). **(H)** Visual comparison of original data to images after SE and segmentation using 3D rendering.

Therefore, we compared the manual and FWHM derived estimates of vascular diameter to those derived from GF and SE after thresholding. No significant differences were found between FWHM, GF, or SE based diameter measurements and their corresponding manual measurements in any of the four vessels. However, GF delivered more variable results with a particular issue when enhancing the PMBC (**Fig. 4C-F**), while SE delivered more consistent results (**Fig. 4G**). These data, together with visual assessment (**Fig. 4G-H**; **Videos 1** and **2**), suggested that neither segmentation method introduced a significant artificial bias and that true biological diameters would be delivered.

**Video 1. Original image 3D rendered**. Video shows original 3D rendered 3dpf *Tg(kdrl:HRAS-mCherry)*^*s916*^.

**Video 2. Image 3D rendered after SE and segmentation**. Video shows 3D rendered 3dpf *Tg(kdrl:HRAS-mCherry)*^*s916*^ after SE and segmentation.

### 3.3. Segmentation Robustness to Noise

Segmentation *robustness* was assessed by processing of data with varying signal properties. To achieve this we examined two approaches, **(a)** acquisition of data with a controlled decrease of image quality (**Fig. 5A-C**) and **(b)** data augmentation by artificial noise addition (Fig. 5D-E), both followed by segmentation after GF and SE. Quantification of the cerebral vascular volume as a segmentation out-read showed no significant difference after GF or SE over the range of tested image qualities (**Fig. 5F;** p 0.3248 and p 0.9981, respectively), suggesting that both segmentation approaches were robust over a broad range of CNR levels. However, despite there being no significant difference in cerebral vascular volume, the coefficient of variation was significantly greater in GF-based measurements (CoV: LP 1.2 18.76%, LP 0.8 14.63%, LP 0.4 17.56%) compared to TF (CoV: LP 1.2 7.81%, LP 0.8 6.97%, LP 0.4 9.14%) for all CNRs, suggesting that while there is no significant difference in accuracy, the GF approach is less precise and consistently so across the range of CNR tested. To test this further, and whether data augmentation would be a suitable replacement to changing data acquisition, artificial noise was added to data and vascular volumes again quantified. CNR was successfully decreased with this approach (**Fig. 5G**) and vascular volume was statistically significantly increased with GF (p 0.0247) but not TF (p>0.9999). Similar to experimentally derived data, GF CoV (31.56%) was larger than TF CoV (9.69%) in augmented data.

**Figure 5.**
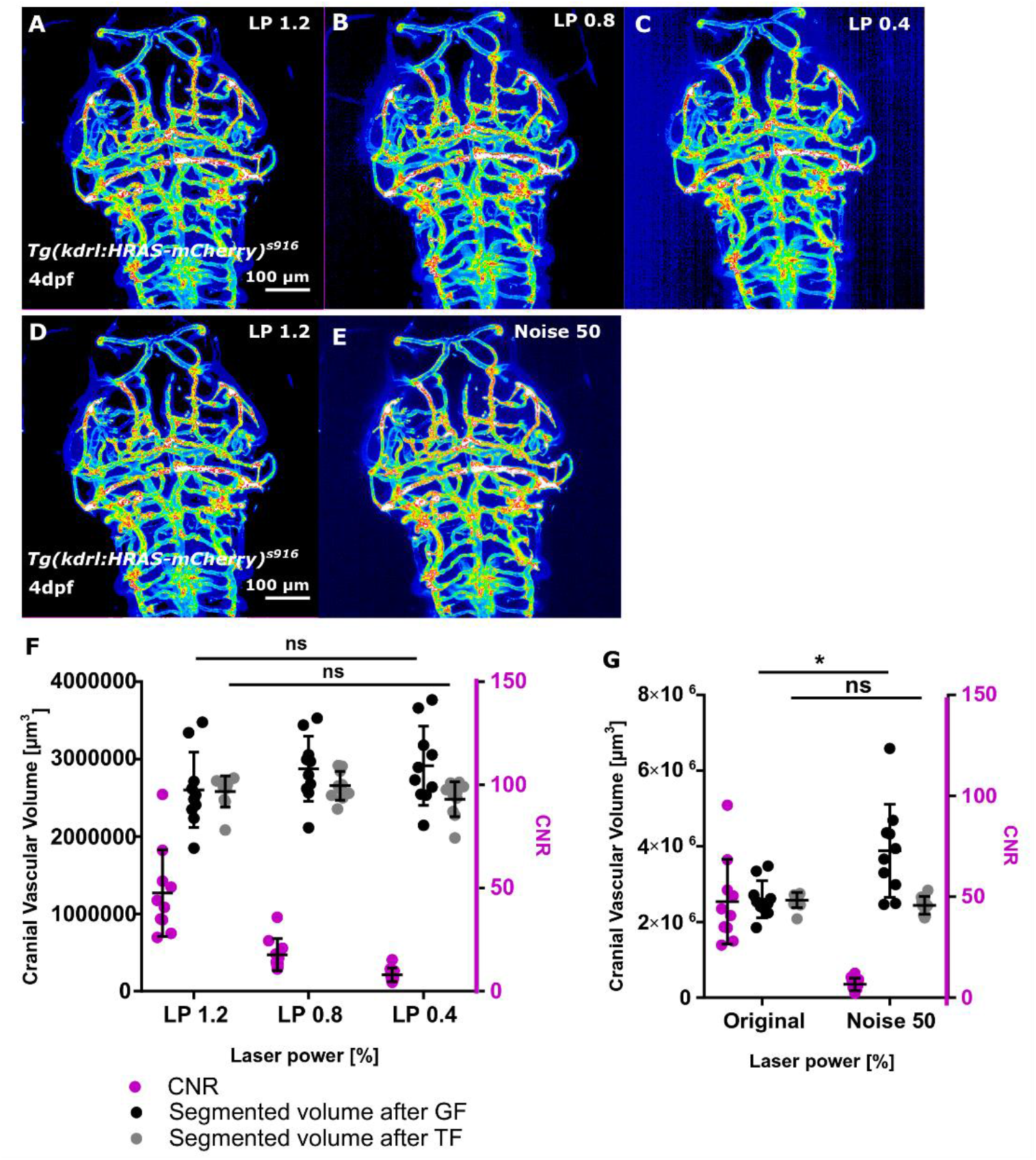
Validation of segmentation robustness. **(A-C)** Dataset with decreased image quality was produced by repeated image acquisition with reduced laser power (LP; 1.2%, 0.8% and 0.4%). **(D-E)** Data augmentation was achieved by addition of noise to images of LP 1.2. **(F)** In experimentally derived data, CNR (magenta) was decreased with LP decrease. No statistically significant difference of vascular volume was observed after GF (p 0.3248; black) or SE (p 0.9981; grey) by LP reduction (n=10 4dpf embryos; 2 experimental repeats; One-Way ANOVA). **(G)** In augmented data, CNR was also decreased (magenta). Vascular volume was statistically significantly increased following GF (p 0.0247) but not TF (p>0.9999).

### 3.4. Segmentation Sensitivity to Biological Differences

To test segmentation *sensitivity*, datasets with a predictable and known biological vascular volume difference were acquired before and after exsanguination (**Fig. 6A,B**). Quantification of the cerebral vascular volume after GF showed no statistically significant difference between control and exsanguinated samples (p 0.2596; **Fig. 6C**; mean value difference 7.8%), while a statistically significant reduction was found after SE (p<0.0001; **Fig. 6D**; mean value difference 8.05%). Importantly, CoV after GF was found to be 38.26% and 26.28% in controls and exsanguinated samples respectively, while CoV was only 10.22% and 9.65% in controls and exsanguinated samples after SE. To confirm that data quality was not significantly altered by sample removal for the exsanguination procedure, CNR was quantified in the basilar artery (BA) before and after exsanguination and no statistically significant difference found (p 0.0876; **Fig. 6E**), indicating that data quality was largely unaltered. Furthermore, the CNR measured in both cases was towards the upper end of the range over which the methods have been shown to be accurate. These data indicated that SE was more sensitive, allowing the extraction of true biological differences due to the intrinsic lower variability of the method. In summary, all of the experiments performed so far to assess the accuracy, robustness and sensitivity of the segmentation approaches indicated that SE-based segmentation is more successful than the GF-based approach.

**Figure 6.**
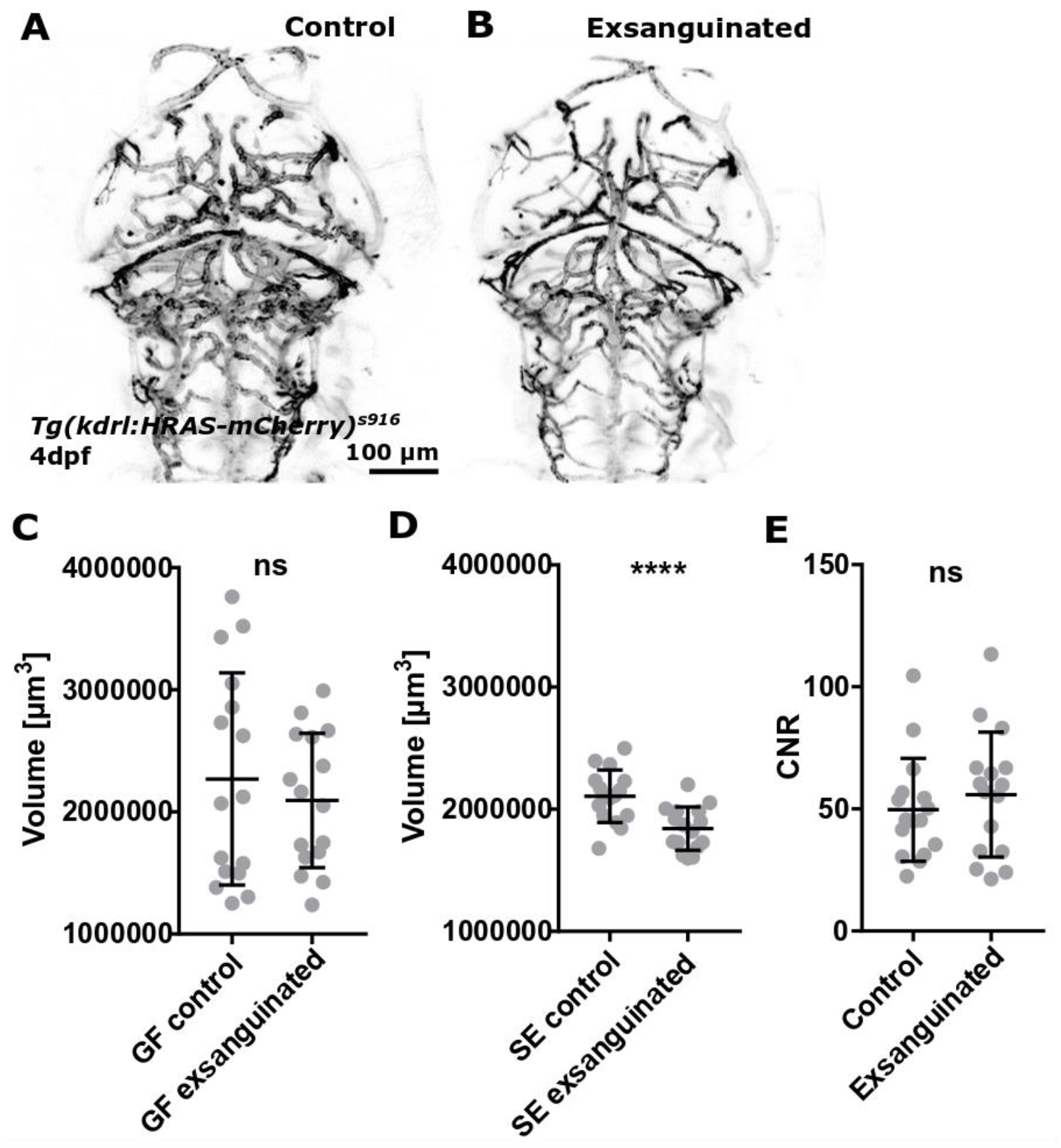
Validation of segmentation sensitivity. **(A-B)** Data were acquired before and after exsanguination. **(C)** Vascular volume quantification after GF showed no statistically significant difference between control and exsanguinated samples (p 0.2596; n=16 4dpf embryos; 2 experimental repeats; paired t-test). **(D)** Vascular volume quantification after SE showed a statistically significant decrease between control and exsanguinated samples (p<0.0001; paired t-test). **(E)** CNR was not statistically significant changed by the exsanguination procedure (p 0.0876; paired t-test).

### 3.5. Quantification of cerebral vascular volume in development

The promising segmentation results in *Tg(kdrl:HRAS-mCherry)*^*s916*^ using SE combined with Otsu thresholding encouraged us to quantify vascular volume from 3-to-5dpf to study cerebrovascular volume in development. After both GF and SE, a statistically significant increase of vascular volume was observed (p 0.0009 and p<0.0001, respectively; **Fig. 7A,B**), but CoV was again higher after GF (3dpf 17.18%, 4dpf 17.96%, 5dpf 27.14%) than TF (3dpf 12.94%, 4dpf 14.59%, 5dpf 13.20%). Visual assessment showed satisfying segmentation results after SE (**Fig. 7C**). Importantly, these data suggested that SE-processed data were less variable and therefore lower sample numbers would be required to extract biologically relevant vascular volumes. This supports the previous findings presented in Section 3.4, where biological differences were only found when using vascular volumes derived from TF-based segmentation and not from GF. This is also exemplified by the fact that mean vascular volume increases are about 12% (GF 3-4dpf 11.42%, 4-5dpf 12.59%; TF 3-4dpf 11.99%, 4-5dpf 11.39%), thus accurate measurements are needed to extract subtle differences.

**Figure 7.**
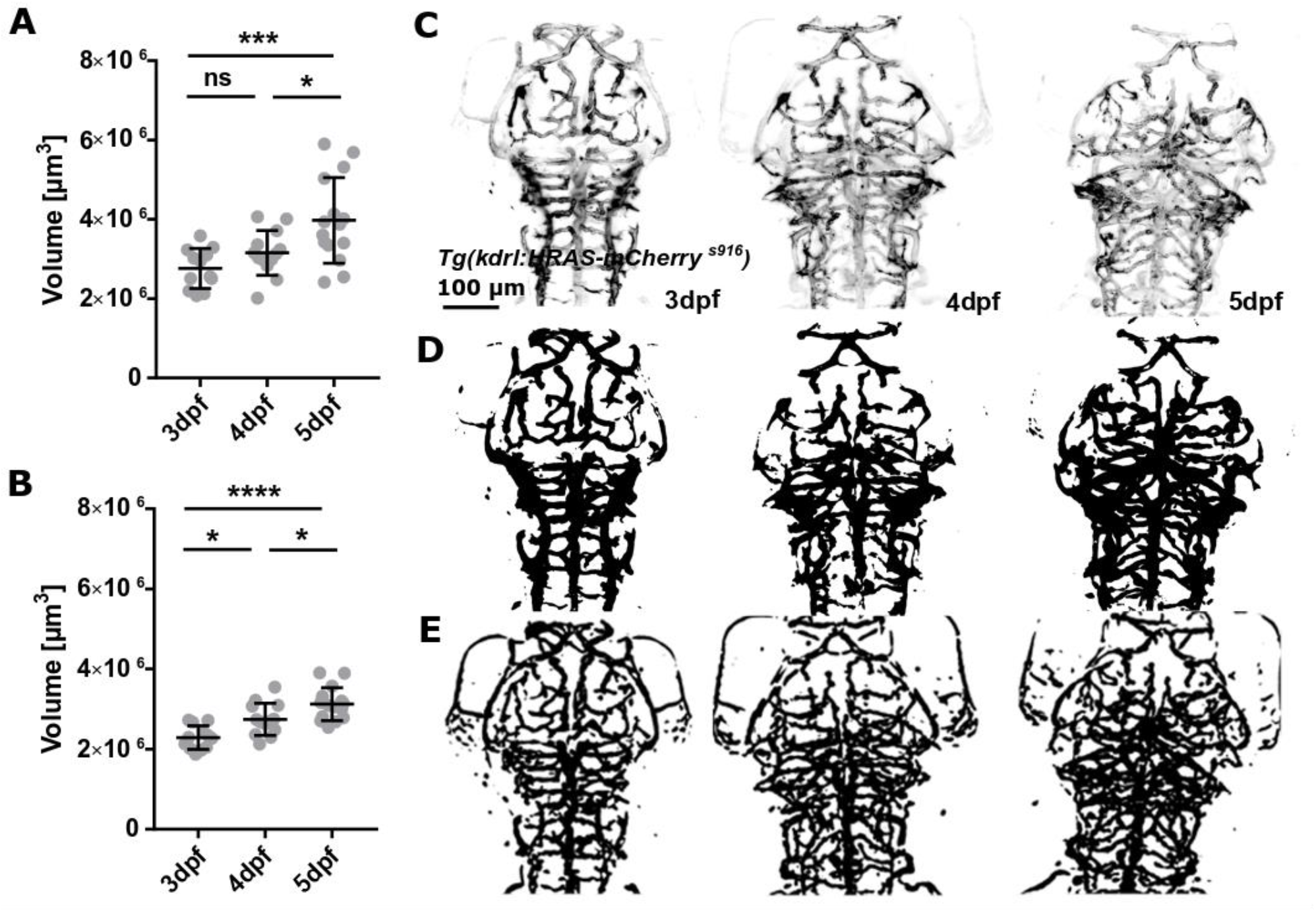
Application to quantify cerebral vascular volume. **(A)** Quantification of vascular volume after GF showed a statistically significant increase from 3-to-5dpf (p 0.0009; 3dpf n=12, 4dpf n=13, 5dpf n=15; 2 experimental repeats; One-Way ANOVA). **(B)** Quantification of vascular volume after SE showed a statistically significant increase from 3-to-5dpf (p<0.0001; One-Way ANOVA). **(C)** Visual comparison of original data with segmented after GF (**D**) and SE (**E**) suggested SE delivered better results.

### 3.6. Application to other transgenic lines

As our enhancement and segmentation approach was optimized for the transgenic reporter line *Tg(kdrl:HRAS-mCherry)*^*s916*^, there was the rational to evaluate whether the suggested approach would be generalizable to other transgenic lines. Thus, we examined the cerebrovascular volume in three different double-transgenics, namely **(1)** *Tg(fli1a:eGFP*)^*y1*^, *Tg(kdrl:HRAS-mCherry*)^*s916*^, **(2)** *Tg(fli1a:CAAX-eGFP), Tg(kdrl:HRAS-mCherry*)^*s916*^, and **(3)** *Tg(fli1a:LifeAct-mClover*)^*sh467*^, *Tg(kdrl:HRAS-mCherry*)^*s916*^. We hypothesised that signal driven under the *fli1a* promotor would be more challenging to segment due to lower vascular specificity and higher image artefact levels, such as reflection from the skin (**Fig. 8A,C**).

**Figure 8.**
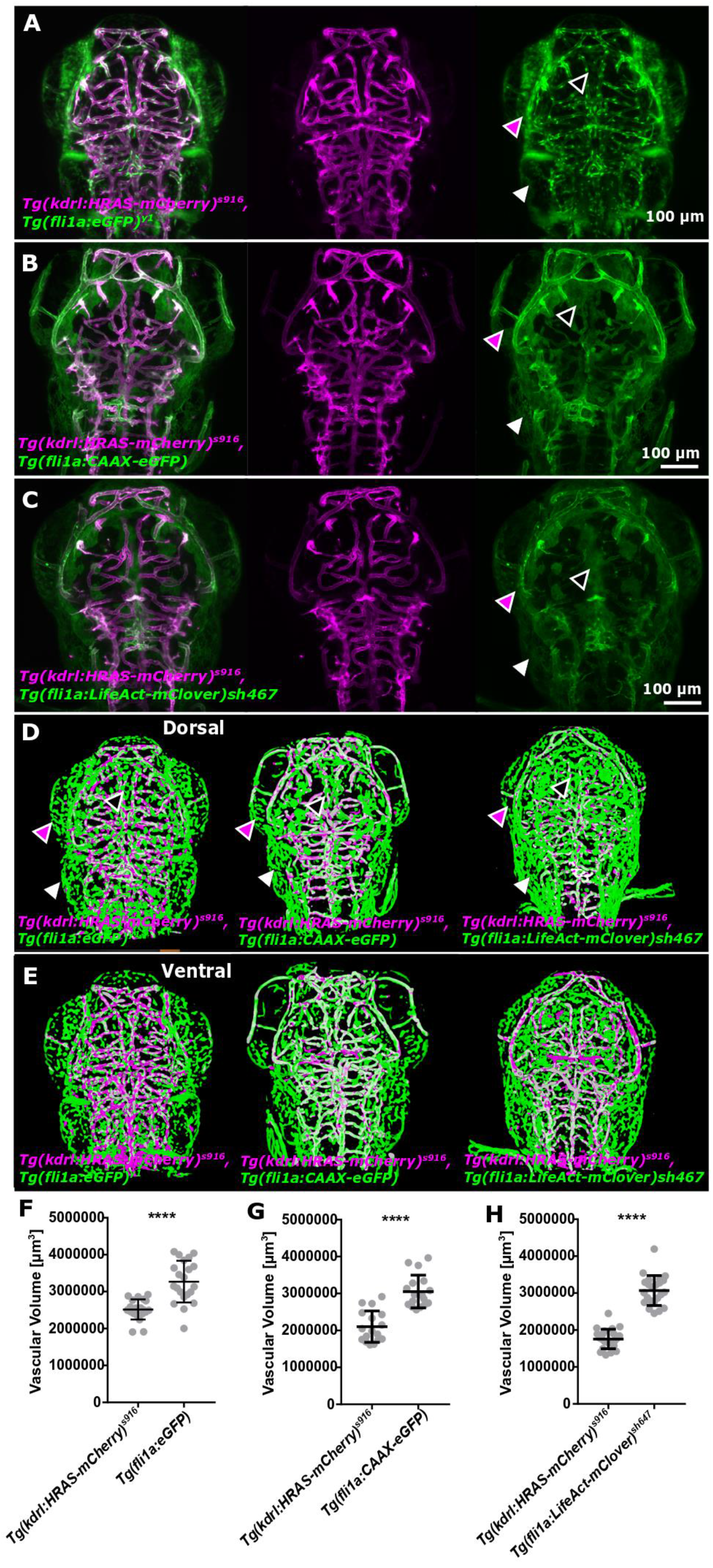
Segmentation of different transgenic lines. **(A-C)** In all three double-transgenics (**(1)** *Tg(fli1a:eGFP*)^*y1*^, *Tg(kdrl:HRAS-mCherry*)^*s916*^, **(2)** *Tg(fli1a:CAAX-eGFP), Tg(kdrl:HRAS-mCherry*)^*s916*^, and **(3)** *Tg(fli1a:LifeAct-mClover*)^*sh467*^, *Tg(kdrl:HRAS-mCherry*)^*s916*^) non-vascular signal was observed in the *fli1a* driven transgenic (arrowheads). **(D-E)** Segmentation results of the three double-transgenics showed non-vascular signal to be enhanced and segmented in the transgenics under *fli1a* promotor. **(F)** Vascular volume in *Tg(fli1a:eGFP*)^*y1*^ was statistically significantly higher than *Tg(kdrl:HRAS-mCherry*)^*s916*^ (p<0.0001; n=21 paired; t-test). **(G)** Vascular volume in *Tg(fli1a:CAAX-eGFP)* was statistically significantly higher than *Tg(kdrl:HRAS-mCherry*)^*s916*^ (p<0.0001; n=17paired; t-test). **(H)** Vascular volume in *Tg(fli1a:LifeAct-mClover*)^*sh467*^ was statistically significantly higher than *Tg(kdrl:HRAS-mCherry*)^*s916*^ (p<0.0001; n=23 paired; t-test).

Segmentation results in the reporter lines under the *fli1a* promotor showed enhancement and segmentation of non-vascular signal (**Fig. 8D-E**).

Quantification of the cerebral vascular volume showed a statistically significantly higher vascular volume under the *fli1a* promotor in all three examined lines (p<0.0001 for all three; **Fig. F-H**). Even though CoV were low (17.28% *Tg(fli1a:eGFP*)^*y1*^, 14.55% *Tg(fli1a:CAAX-eGFP)*, 13.18% *Tg(fli1a:LifeAct-mClover*)^*sh467*^) this exemplifies that further optimization and processing will be needed to reliably extract vascular signal from transgenics other than *Tg(kdrl:HRAS-mCherry*)^*s916*^ for which our approach was optimized.

Together, this suggests that vascular volume can be extracted in other transgenics, but accuracy and precision are reduced. The challenges encountered, showed that non-vascular signal (pan-endothelial) as well as non-specific (eg. skin) signal both lead to an increase in extracted vascular volume. To address this, future work might include pre-processing to enhance vascular and decrease non-vascular signal, exclusion of non-connected components, exclusion of objects under a specified size-threshold, or exclusion based on peripheral position.

### 3.7. Deep Learning

In addition to the conventional image processing approaches, based on GF and SE filtering combined with Otsu thresholding, we aimed to develop the first deep learning approach for segmentation of zebrafish vascular data from the single transgenic *Tg(kdrl:HRAS-mCherry*)^*s916*^. To achieve this, we trained an original U-Net, SegNet, and three modified U-Nets (dU-Net1-3) on our training dataset using segmented masks obtained after SE as the ground truth (**Fig. 9A-C**) and the resulting trained network was then applied to the evaluation dataset.

**Figure 9.**
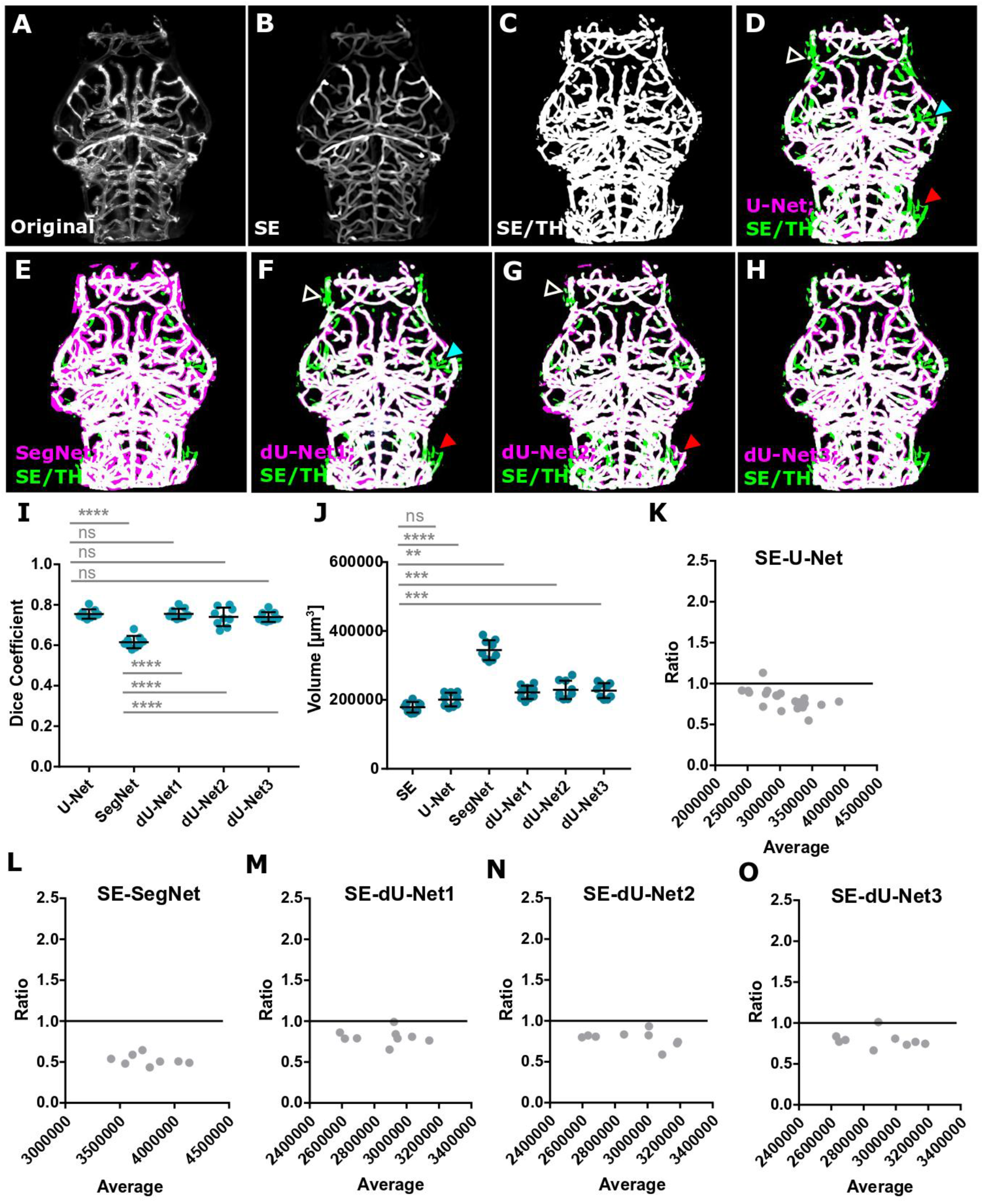
Machine learning results when trained from original data. **(A)** Original image, **(B)** enhanced, **(C)** segmented using Otsu thresholding (referred to as SE/TH). **(D)** MIP of SE/TH (green) and original U-Net (magenta) segmentation, showing high degrees of overlap (white), while certain vessels were extracted with SE but not U-Net (arrowheads). **(E)** MIP of SE/TH (green) and SegNet (magenta) segmentation, showing consistent over-segmentation with SegNet. **(F)** MIP of SE/TH (green) and dU-Net1 (magenta) segmentation, showing high degrees of overlap (white), while certain vessels were extracted with SE/TH but not dU-Net1 (arrowheads). **(G)** MIP of SE/TH (green) and dU-Net2 (magenta) segmentation. **(H)** MIP of SE/TH (green) and dU-Net3 (magenta) segmentation. **(I)** Dice Coefficient of segmentation outcomes. **(J)** Quantified vascular volumes. **(K-O)** Bland-Altman ratio test comparing vascular volume values.

The results of the ML-based segmentation approaches were compared visually to SE-based segmentation **(Fig. 9D-H**; example 3D renderings are provided in **Videos 3-7**). Quantification of the evaluation dataset produced Dice Coefficient (compared to the SE based segmentation) (**Fig. 9I**), vascular volume (**Fig. 9J**), and Bland-Altman ratio (**Fig. 9K-O**) showed U-Net architectures to deliver better results than SegNet, which over-segmented the vasculature (**Table 2**). Comparing the different U-Net architectures, the original U-Net and dU-Net1 delivered the best results with an average Bland-Altman ratio of 0.896 and 0.809 and a vascular volume CoV of 9.78% and 8.65% for U-Net and dU-Net1, respectively. There was a tendency for the deep learning methods to systematically over-estimate the vascular volume and all except the original U-Net showed statistically significant increases. With the exception of the SegNet output, the differences were generally small with comparable CoV to the SE based volume measurements, suggesting that the deep learning methodology should have comparable sensitivity to volume changes, even though those volumes are slightly over-estimated.

**Table 2.**
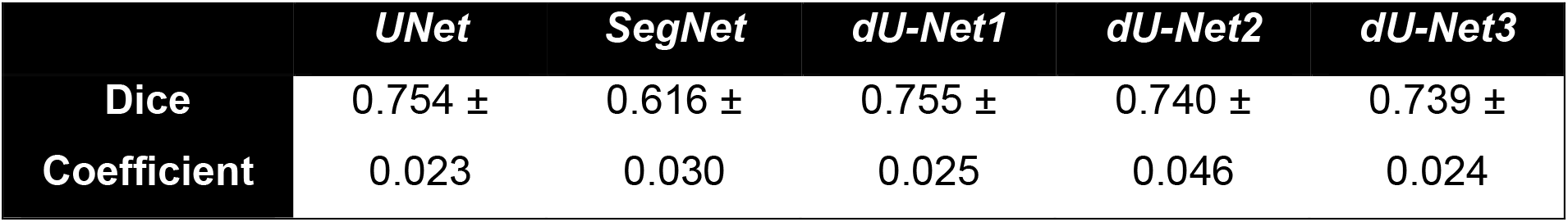
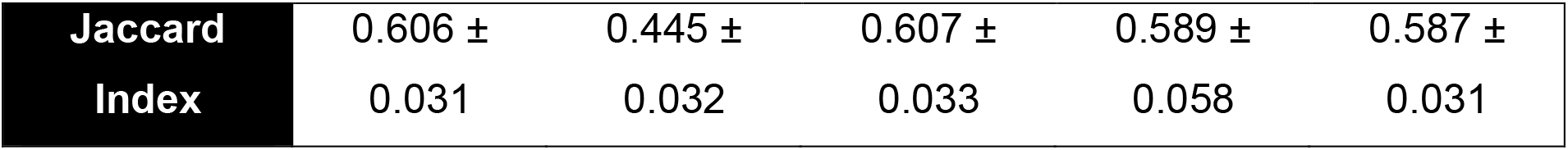
Dice Coefficient and Jaccard Index of ML-based segmentation approaches in comparison to SE-based segmentation.

This is likely due to the fact that SegNet architectures are usually used to segment natural images with sharp edges (such as landscapes), while the original U-Net was developed for biomedical images.

Together, this showed that deep learning based segmentation can be applied to our data, and that U-Net based architectures outperform the SegNet architecture, while changes in the U-Net architecture (convolutional layers, employing batch normalization, and dropout procedures) did not significantly impact segmentation outcomes.

The deep learning approaches presented here operate in 2D on a slice-by-slice basis with a 1-2 second segmentation time (after training) per slice (approx. 400-700 slices per stack) resulting in typical run times of between 7 and 23 minutes for segmenting a full stack. In contrast, SE-based segmentation typically require about 50 minutes in total per stack, suggesting a significant time benefit in favour of the deep learning approach. Nevertheless, future work is needed to examine the performance of deep learning approaches in zebrafish at different ages, in different transgenic lines, exploring and optimising alternative network configurations, implementing a 3D segmentation approach, and investigating alternative methods for generating training data that are independent of our SE-based method.

In summary, we have found that deep learning approaches provide a promising alternative to conventional image processing methods for zebrafish vasculature segmentation, with U-Net based architectures performing particularly well.

**Video 3. 3D rendered segmentation results of U-Net**. Video shows 3D rendered 3dpf *Tg(kdrl:HRAS-mCherry*)^*s916*^ after SE-based segmentation (green) and U-Net segmentation (magenta).

**Video 4. 3D rendered segmentation results of SegNet**. Video shows 3D rendered 3dpf *Tg(kdrl:HRAS-mCherry*)^*s916*^ after SE-based segmentation (green) and SegNet segmentation (magenta).

**Video 5. 3D rendered segmentation results of dU-Net1**. Video shows 3D rendered 3dpf *Tg(kdrl:HRAS-mCherry*)^*s916*^ after SE-based segmentation (green) and dU-Net1 segmentation (magenta).

**Video 6. 3D rendered segmentation results of dU-Net2**. Video shows 3D rendered 3dpf *Tg(kdrl:HRAS-mCherry*)^*s916*^ after SE-based segmentation (green) and dU-Net2 segmentation (magenta).

**Video 7. 3D rendered segmentation results of dU-Net3**. Video shows 3D rendered 3dpf *Tg(kdrl:HRAS-mCherry*)^*s916*^ after SE-based segmentation (green) and dU-Net3 segmentation (magenta).

## 4. Conclusion

In this work, we demonstrated that enhancement and segmentation of the zebrafish vasculature is possible using a variety of approaches based on general filtering, Sato enhancement, or deep learning. Validation of GF- and SE-based approaches involved studying their robustness, sensitivity, as well as applicability to other transgenics. In all cases, we found that the SE-based segmentation outperformed results based on GF. We successfully quantified cerebrovascular volume from 3-to-5dpf, indicating that biologically relevant vascular volumes can be objectively quantified. Once validated, we used SE-based segmentation to train deep learning methods, finding that U-Net based architectures returned vascular segmentation with comparable accuracy and precision.

In conclusion, the proposed segmentation allows for quantification of cerebral vascular volume, facilitating the study of mechanisms of vascular development and disease, as well as the effect of drugs or chemical components. Importantly, the application of robust objective quantification allows for the reduction of sample size needed to assess the vascular phenotype, which is important from an ethical as well as computational point of view.

## Abbreviations

BA: basilar artery
CNR: contrast-to-noise ratio
CtA: central artery
dpf: days post fertilisation
FWHM: Full-Width-Half-Maximum
GF: General Filtering
hpf: hours post fertilisation
LSFM: light sheet fluorescence microscopy
MMCtA: middle mesencephalic central artery
PMBC: primordial midbrain channel
ROI: region of interest
SE: Sato Enhancement

## Acknowledgements

The authors are grateful to the funders for supporting this project, including the Department of Infection, Immunity and Cardiovascular Disease (The University of Sheffield, UK), the Insigneo Institute for *in silico* Medicine, and the British Heart Foundation. The authors would like to thank Professor Holger Gerhardt, Dr Robert Wilkinson, and Dr Aaron Savage for provision of transgenic lines.

